# Electron Transport Chain-inhibiting fungicides increase resistance to widely used azole antifungals

**DOI:** 10.64898/2026.11.19.767815

**Authors:** Severina M. Pociunaite, Sharon A. White, Pin Tong, Rebecca Yeboah, Alison L. Pidoux, Robin C. Allshire

**Author notes:** These authors contributed equally (Severina M. Pociunaite, Sharon A. White).

## Abstract

Rising antifungal resistance in human and crop fungal pathogens poses significant challenges. Azole-based antifungals that inhibit ergosterol biosynthesis are a key first-line treatment for human fungal infections. Importantly, azole-based ergosterol-inhibiting fungicides are also widely used in agriculture alongside electron transport chain (ETC) Complex III inhibiting compounds. However, it is unknown whether the combined use of these distinct fungicides is entirely synergistic. We find that exposure of fission yeast, *Schizosaccharomyces pombe* to ETC-Complex III inhibitors, prior to or simultaneously with azoles, attenuates azole efficacy by inducing mitochondrial dysfunction. Our analyses show that reactive oxygen species (ROS) accumulation as a result of ETC inhibition activates oxidative stress response pathways thereby decreasing azole potency. Related responses in pathogenic fungi indicate that the use of ETC inhibiting fungicides in agriculture may inadvertently promote azole resistance. We conclude that the use of specific fungicide combinations should be re-evaluated in order to preserve both their clinical and agricultural effectiveness.

## Introduction

Fungal pathogens are a major health threat [1, 2], exacerbated by rising antifungal resistance and the availability of only four classes of antifungal drug: azoles, polyenes, echinocandins and allylamines [3–5]. Resistance to azoles, the frontline treatment for human fungal infections, has increased dramatically over the last twenty-five years [5–7]. Azole-based antifungals inhibit Erg11/Cyp51 lanosterol 14*α*-demethylase (commonly known as <u>D</u>e<u>M</u>ethylase Inhibitors or DMIs in agriculture) and are widely used to combat human and crop fungal pathogens [4–8]. Azole-based antifungals, such as Fluconazole (FLC^DMI^), remain the only class of orally administered human medication that is frequently prescribed alone, particularly in resource-limited settings, to ameliorate human *Cryptococcus neoformans* and other fungal infections [9, 10].

In agriculture azoles^DMI^ are used combinatorially to combat plant fungal pathogens that threaten cereal crop yield and have substantial economic impacts [7, 8, 11]. For example, to tackle the pathogen *Zymoseptoria tritici*, winter wheat is sprayed up to four times per season with azoles^DMI^ alongside assorted compounds targeting the mitochondrial electron transport chain (ETC), such as ETC-Complex II Succinate Dehydrogenase Inhibitors (SDHIs) and ETC-Complex III cytochrome bc1 inhibitors (Quinone inside Inhibitors; QiIs and Quinone outside Inhibitors; QoIs) [12–16]. Consequently, residual levels of these compounds can remain in soil and plant stubble, potentially selecting for resistance in the target crop fungal pathogen and any resident environmental fungi including potential human or animal fungal pathogens [8, 17].

*C. neoformans* has been isolated from plant material across a wide range of plant species and geographical locations, suggesting that it commonly inhabits plant-associated environments as either an epiphyte or endophyte. This ecological association may also explain its frequent isolation from pigeon guano, as birds can acquire the fungus through ingestion of colonised plant material and associated insects, subsequently disseminating it in their droppings [18]. Thus, this opportunistic human fungal pathogen can be distributed over ecosystems by bird movement [19, 20]. Hence, there is likely to be a reservoir of *C. neoformans* in crop fields that can be exposed to ETC inhibitor treatments. Currently used ETC-complex inhibitors and azoles^DMI^ have field half-lives ranging from 4 to more than 250 days [21]; the degree to which these compounds persist in the environment is dependent on a myriad of factors. Nevertheless, it is apparent that innocuous environmental fungi, such as *C. neoformans* within the soil or plant matter can be exposed to pockets of residual low levels of ETC-inhibitors persisting in the environment. This exposure has the potential to increase the population of transiently resistant fungal cells growing in the presence of sublethal azole^DMI^ concentrations which may then create a window of opportunity that can lead to the development of stable genetic resistant mutations.

Therefore, although agricultural fungicides have distinct targets, their combined use could trigger unexpected consequences such as suppression, a well-known issue with combinatorial antibiotic use [22]. Mitochondrial dysfunction has been shown to confer resistance to azoles^DMI^ in the non-pathogenic budding yeast (*Saccharomyces cerevisiae)* and fission yeast (*Schizosaccharomyces pombe)* and in several pathogenic fungi [23–27]. Here we explore if and how exposure to combinations of azoles^DMI^ and mitochondrial ETC inhibitors alters azole^DMI^ effectiveness in *S. pombe*, the human pathogen *C. neoformans* and wheat pathogen *Z. tritici*.

## Results

### Mutation of either azole^DMI^ or QiI targets confers resistance to clinical and agricultural azoles^DMI^

Both azole^DMI^-based clinical and agricultural antifungals target the active site of Erg11/Cyp51 and mutations in this enzyme confers azole^DMI^ resistance in many fungi such as *Candida auris* and *Candida parapsilosis (erg11^Y132F^*), *Aspergillus fumigatus (erg11^Y121F^*) as well as in *C. neoformans (CYP51^Y145F^)* and *Z. tritici (CYP51^Y137F^)* [17, 28–31]. To determine if an equivalent mutation in fission yeast confers resistance to both clinical and agricultural azoles^DMI^ we introduced a missense mutation (Y117F) at the endogenous *erg11* locus. The resulting *erg11^Y117F^*cells showed increased resistance to clinical (FLC^DMI^) and agritech (prothioconazole-desthio, PRD^DMI^) azole antifungals compared to wild-type (**Fig 1A-1C**), indicating that any selection imposed by agritech azole^DMI^ fungicides could drive resistance to clinical azole^DMI^-based antifungals in environmentally located opportunistic fungi that may subsequently infect humans.

**Figure 1.**
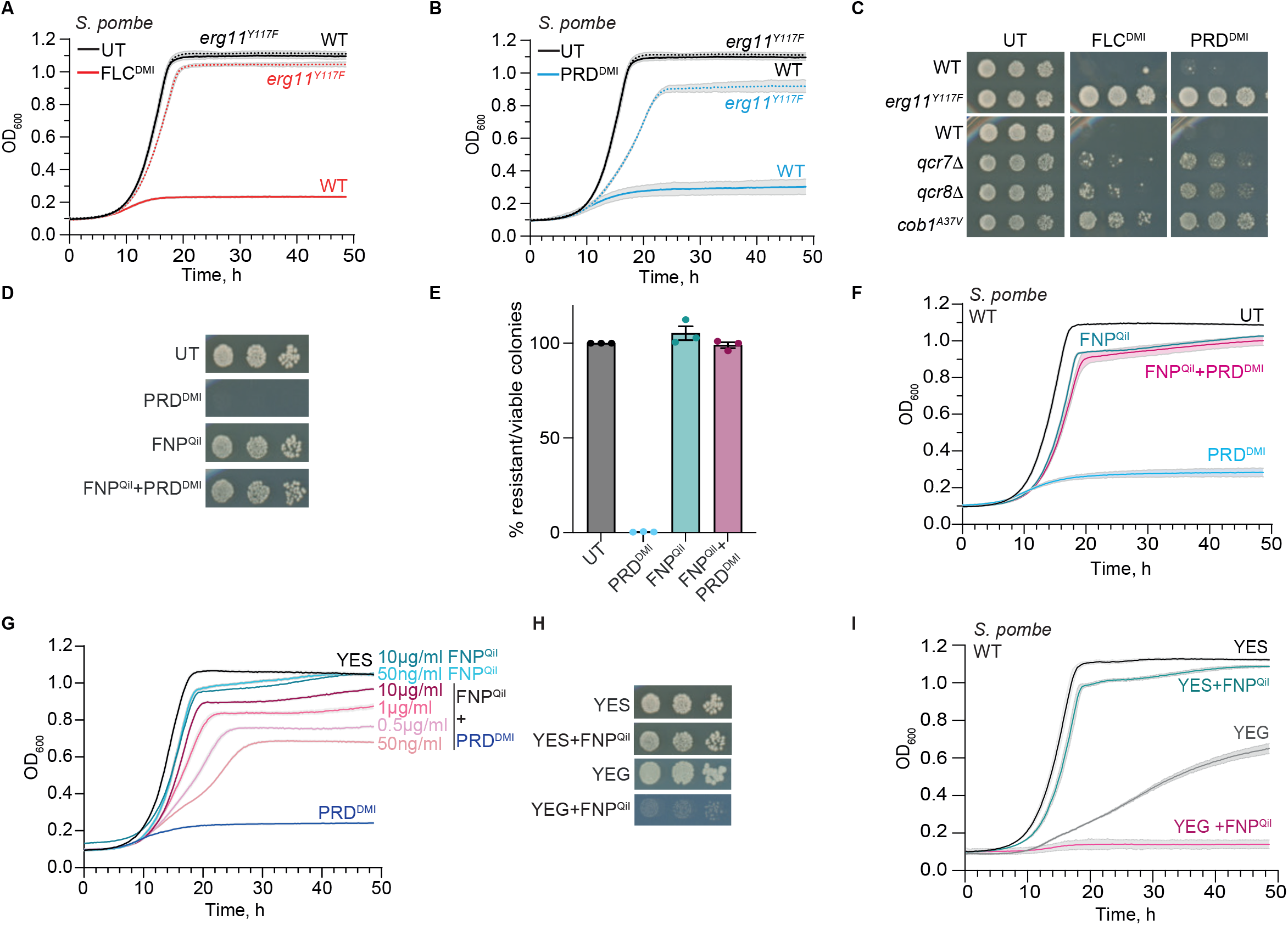
FNP^QiI^ fungicide renders fission yeast resistant to azole^DMI^. (A) WT (solid lines) and *erg11^Y117F^*(dashed lines) *S. pombe* cells grown in YES (UT, untreated, black) or fluconazole (FLC^DMI^, red). Growth curves: average three replicates. Shading: variation. (B) Growth curves (as A) grown with/without prothioconazole-desthio (PRD^DMI^, blue). (C) Comparative growth assay comparing WT to *erg11^Y117F^* (top), or *qcr7Δ, qcr8Δ,* and *cob1^A37V^* (bottom) on YES alone (UT), or with addition of FLC^DMI^ or PRD^DMI^. (D-F) Comparative growth assays of WT cells on YES alone (UT), or with addition of PRD^DMI^, FNP^QiI^ or PRD^DMI^+FNP^QiI^. (E) Percentage colony forming units. (F) Growth curves as (A). *S. pombe* cells grown in YES (UT, untreated, black), prothioconazole-desthio (PRD^DMI^, blue), fenpicoxamide-phenol (FNP^QiI^, teal) or both PRD^DMI^+FNP^QiI^ (dark pink). (G) Growth curves of WT cells in YES alone (UT), or with addition of FNP^QiI^, PRD^DMI^, or FNP^QiI^+PRD^DMI^, with concentrations of FNP^QiI^ varying as indicated. (H) Comparative growth assays on YES (glucose) or non-fermentable YEG (glycerol) plates +/-FNP^QiI^. (I) Growth curves: WT cells in YES +/-FNP^QiI^ compared to YEG +/-FNP^QiI^.

ETC-Complex III inhibitors such as QiIs target the mitochondrially-encoded Cob1 (cytochrome b) subunit of the ubiquinol-cytochrome-c reductase complex. It is currently not possible to target mutations to specific locations of the fission yeast mitochondrial genome, hence we utilized a previously isolated Cob1 mutant, *cob1^A37V^*[32]*. S. pombe* cells harboring the *cob1^A37V^*active site mutation are resistant to the ETC complex III inhibitor antimycin A (ANT^QiI^) but also to FLC^DMI^ and PRD^DMI^ [32] (**Fig 1C**, S1A and S1B Fig). Similarly, cells lacking Qcr7 or Qcr8 ETC-Complex III components exhibited resistance to both of these azoles (*qcr7*Δ and *qcr8*Δ; **Fig 1C**, S1C and S1D Fig), as do other mutations and epimutations impairing mitochondrial function [23, 24, 32]. Thus, azole^DMI^ resistance in fission yeast can arise from direct mutation of the Erg11 target, or indirectly via impaired function of the ETC-Complex III/ubiquinol-cytochrome c reductase complex.

### Treatment with the QiI fenpicoxamid-phenol^QiI^ suppresses azole^DMI^ sensitivity in fission yeast

Distinct QiI and QoI fungicides that inhibit ETC-Complex III function are used in agriculture in combination with azoles to protect cereal crops from fungal infections [12]. The QiI antimycin A (ANT^QiI^) is primarily used as a pesticide but QiI fungicides, such as fenpicoxamid-phenol (FNP^QiI^), were recently introduced particularly to control Septoria Tritici/Leaf Blotch in wheat caused by *Zymoseptoria tritici* [14–16]. In contrast QoIs (also called strobilurins) such as azoxystrobin (AZS^QoI^) have been used to protect a range of crops for almost 30 years and are still used in crop protection, although they are no longer effective against *Z. tritici* [13, 14, 33].

Since ETC-Complex III mutations conferred resistance to azole^DMI^ antifungals in *S. pombe* (**Fig 1C** and S1A-S1D Fig) we predicted that ETC-Complex III inhibition might similarly induce resistance. Exposure of wild-type *S. pombe* cells to the ETC-Complex III inhibitor FNP^QiI^ had little impact on growth in glucose-rich media (YES), which favors fermentation rather than mitochondrial-based respiration to produce ATP [34, 35] (**Fig 1D-1F**). Nevertheless, wild-type *S. pombe* sensitivity to PRD^DMI^ and FLC^DMI^ was suppressed by addition of FNP^QiI^, rendering cells PRD^DMI^ and FLC^DMI^ resistant (**Fig 1D-1F** and S1E Fig). This suppression of PRD^DMI^ and FLC^DMI^ sensitivity is also dependent on FNP^QiI^ concentration (**Fig 1G**). Moreover, FNP^QiI^ treatment synergized with the ETC-complex III *cob1^A37V^*and *qcr7Δ* mutants to confer even greater PRD^DMI^ resistance (S1F and S1G Fig). Consistent with ETC-Complex III inhibition by FNP^QiI^, its addition curtailed *S. pombe* growth in glycerol, a non-fermentable carbon source, which forces ETC-mediated respiration for ATP production [34, 35] (**Fig 1H** and **1I**).

Addition of the ETC-Complex III inhibitors AZS^QoI^ or ANT^QiI^ also increased *S. pombe* resistance to FLC^DMI^ and PRD^DMI^ azoles (**Fig 2A-C**), and further increased FLC^DMI^ and PRD^DMI^ resistance of *cob1^A37V^*and *qcr7*Δ mutants (**Fig 2C**). Moreover, growth inhibition with mefentrifluconazole (MFZ^DMI^), a recently introduced azole^DMI^ fungicide with reported superior efficacy for tackling fungal disease in crops [36] was also suppressed in the presence of FNP^QiI^, ANT^QiI^ or AZS^QoI^, or ETC-Compex III mutations (**Fig 2D and 2E**). To assess whether these effects were specific we also tested thiabendazole (TBZ; microtubule disruptor [37]), a fungicide used primarily as a post-harvest treatment to prevent mould, and KCl that causes osmotic and salt stress as a non-fungicidal stress control. Neither treatment increased resistance of *S. pombe* to the presence of PRD^DMI^ (S1H and S1I Fig). These findings indicate that the observed suppression of azole sensitivity is specific to the combination of complex III QiI/QoIs demonstrating that QiI/QoI treatment selectively compromises azole effectiveness. Thus, exposure of non-pathogenic wild-type fission yeast [38] to ETC-complex III inhibitors that are used to control crop fungal infections, specifically compromises azole^DMI^ antifungal effectiveness. This previously unrecognised interaction between antifungal compounds with distinct modes of action exposes a potential weakness in current agricultural fungal disease-management strategies.

**Figure 2.**
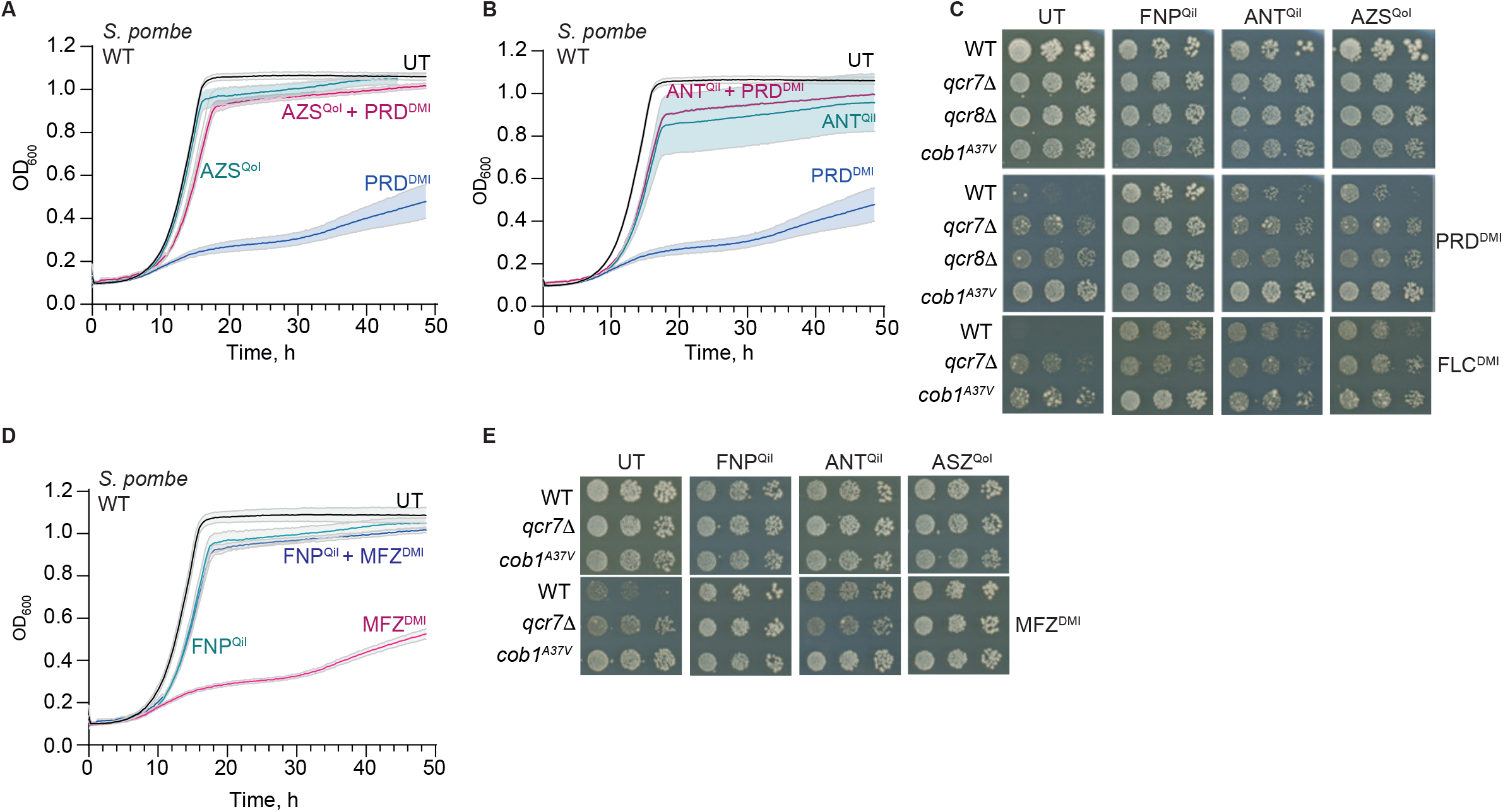
ETC-Complex III Qi and Qo inhibiting fungicides increase azole^DMI^ resistance of *S. pombe*. (A) Growth curves of WT *S. pombe* cells grown in YES (UT, untreated, black), PRD^DMI^ (blue) azoxystrobin (AZS^QoI^, teal), or both PRD^DMI^+ AZS^QoI^ (dark pink). **(B)** Growth curves as (A). *S. pombe* cells grown in YES (UT, untreated, black), PRD^DMI^ (blue) antimycin A (ANT^QiI^, teal), or both PRD^DMI^+ ANT^QiI^ (dark pink). (C) Comparative growth assays of WT, *qcr7Δ, qcr8Δ* and *cob1^A37V^* cells on indicated media arranged in a grid pattern to show effects of drug combinations with PRD^DMI^ or FLC^DMI^. (D) Growth curves of WT cells in YES alone (UT), or with addition of FNP^QiI^ (teal), mefentrifluconazole (MFZ^DMI^, dark pink) or FNP^QiI^+MFZ^DMI^ (blue). (E) Comparative growth assays of WT, *qcr7Δ* and *cob1^A37V^* cells on indicated media arranged in a grid pattern to show effects of drug combinations with MFZ^DMI^.

### FNP^QiI^ induces mitochondrial dysfunction and intracellular ROS

To determine how FNP^QiI^ promotes azole resistance, we assessed its impact on mitochondrial function and downstream consequences. Respiring cells normally reduce the colourless redox indicator TTC (2,3,5-triphenylte-trazolium chloride) to red-coloured TPF (2,3,5-triphenyltetrazolium formazan) [24, 39]. Consequently *S. pombe* ETC-complex III mutants *qcr7*Δ, *qcr8*Δ and *cob1^A37V^* fail to produce red TPF, indicating defective respiration (**Fig 3A**) [24]. Wild-type cells grown on FNP^QiI^-containing plates also failed to convert TTC to red TPF (**Fig 3A**). Thus, impairment of ETC-Complex III function either by genetic mutations or FNP^QiI^ inhibition, promotes mitochondrial dysfunction and PRD^DMI^ resistance.

**Figure 3.**
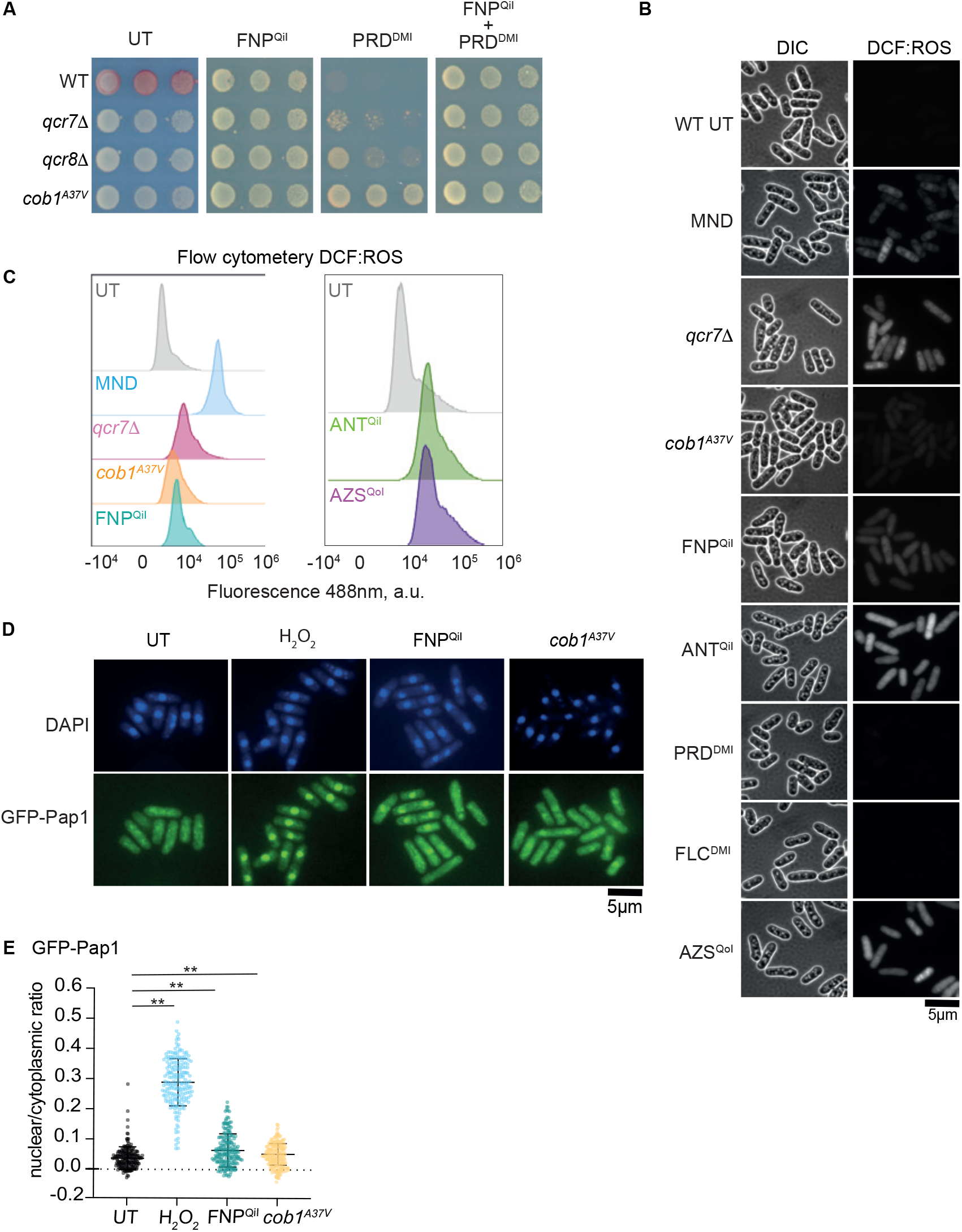
FNP^QiI^ induces mitochondrial dysfunction and intracellular ROS. (A) Respiratory competence TTC overlay assay. Serial dilutions of indicated cells grown on YES alone (UT), or with addition of PRD^DMI^ or FNP^QiI^, or FNP^QiI^+PRD^DMI^. Red indicates respiration. (B) DCF ROS detection. WT cells untreated (UT) or treated with ROS-generator MND, or FNP^QiI^, ANT^QiI^, PRD^DMI^, FLC^DMI^ or AZS^QoI^ in YES or *qcr7Δ* or *cob1^A37V^* mutant cells incubated in DCFH-DA ROS-indicator. Brightfield (left) and 488 nm (right). (C) 488nm flow cytometry profiles of cells (as B) measuring ROS levels by DCF detection. (D, E) GFP-Pap1 localization and quantification (E) in WT cells untreated (UT), treated with H_2_O_2_ (control) or FNP^QiI^, or *cob1^A37V^*mutant cells. n> 200 cells. Significance: Student’s *t*-test; *\*\*p<0.01*.

ETC inhibition is known to result in reactive oxygen species (ROS) accumulation, which can damage ETC components, culminating in ROS leakage into the cytoplasm [40, 41]. Intracellular ROS is detectable via microscopy or flow cytometry using the sensor DCFH-DA which ROS converts to fluorescent DCF [42]. As expected, intracellular ROS levels increased upon treatment of wild-type *S. pombe* cells with the ROS-generator menadione (MND, **Fig 3B and 3C**). Higher intracellular ROS levels were also detected in *qcr7*Δ, *qcr8*Δ and *cob1^A37V^* mutant cells [24]. Importantly, all ETC-Complex III inhibitors (FNP^QiI^, ANT^QiI^ or AZS^QoI^) also increased ROS levels in wild-type cells, whereas PRD^DMI^ and FLC^DMI^ did not. Thus, ROS levels in fission yeast are increased by impairment of ETC-Complex III function through either mutation or exposure to inhibitory fungicides.

### Prior exposure to FNP^QiI^ transiently increases PRD^DMI^ resistance, induces stress responses and increases efflux

In many fungi including yeasts, intracellular ROS induces oxidative stress responses via oxidation sensitive transcription factors such as Pap1 in *S. pombe* or Yap1 in *S. cerevisiae* [43–46]. Oxidative stress causes Pap1 to accumulate in *S. pombe* nuclei where it induces core oxidative stress genes including those encoding oxidoreductases (Grx1, Obr1), sulfiredoxin (Srx1) and efflux pumps/plasma membrane transporters (Bfr1, Caf5, Pmd1) [43, 44, 46]. Since FNP^QiI^ treatment increased intracellular ROS levels, we tested if it induced the oxidative stress response. Increased nuclear accumulation of GFP-Pap1 was detected following hydrogen peroxide (H_2_O_2_) or FNP^QiI^ addition to wild-type cells or in *cob1^A37V^* mutant cells, indicating that FNP^QiI^ treatment may also induce a Pap1-mediated transcriptional response (**Fig 3D and 3E**).

To explore the longevity of FNP^QiI^-stimulated PRD^DMI^ resistance and gene expression changes we followed the dynamics of PRD^DMI^ resistance after FNP^QiI^ treatment. Wild-type *S. pombe* cells were exposed to FNP^QiI^ for 24 hours and subsequently harvested at intervals after FNP^QiI^ removal (**Fig 4A**). Immediately following FNP^QiI^ washout (0 hr) cells displayed increased growth in the presence of PRD^DMI^ which persisted for up to 24 hours following FNP^QiI^ removal, before returning to untreated control levels (**Fig 4B**). Thus, the observed FNP^QiI^-stimulated resistance to PRD^DMI^ is a transient response and likely reflects induction of stress sensing pathways that act to protect cells from perceived environmental threats.

**Figure 4.**
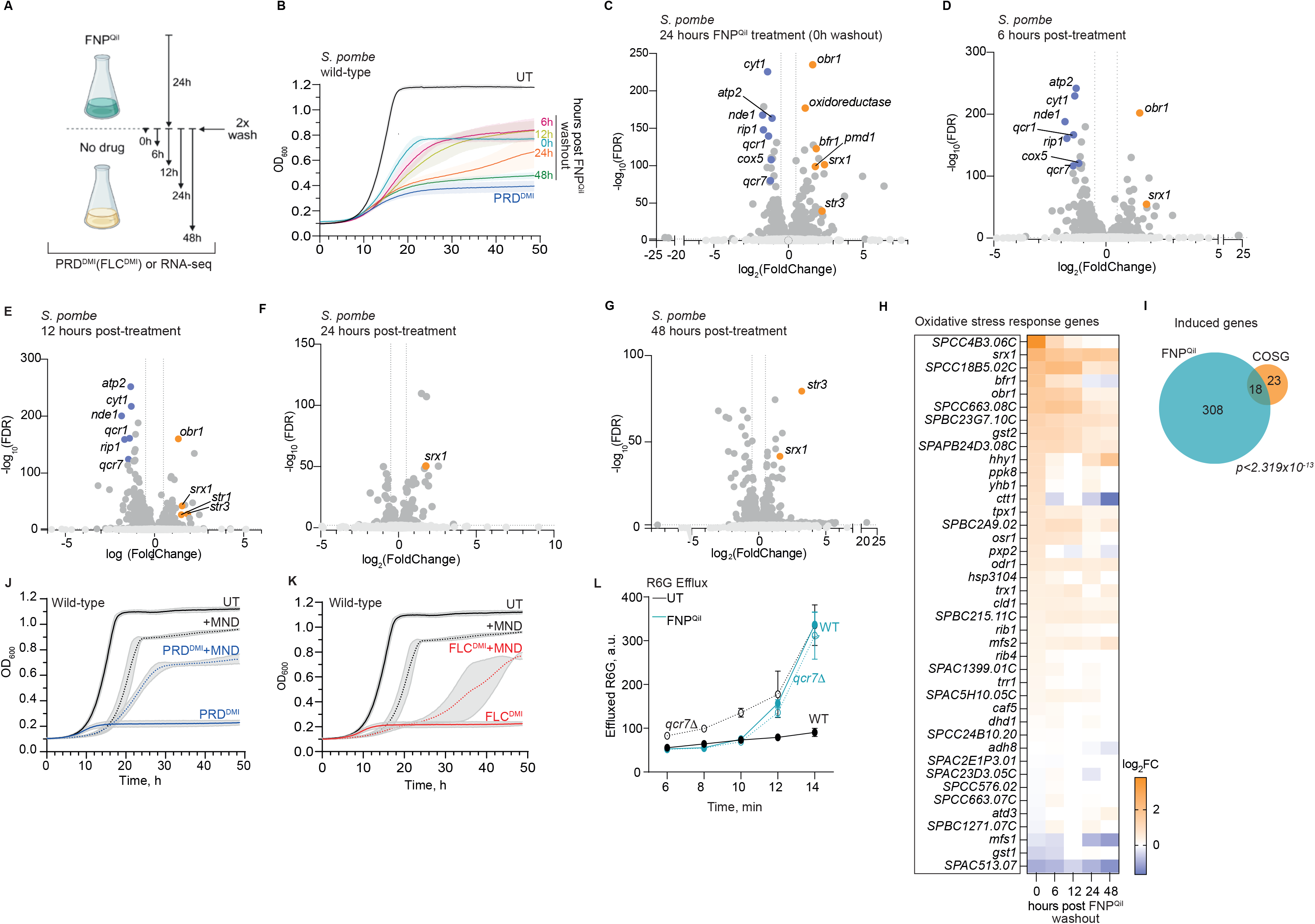
Transient FNP^QiI^ exposure activates the fission yeast Pap1-mediated oxidative stress response resulting in increased efflux. (A) Schematic of experimental set-up. WT *S. pombe* cells are pre-exposed to FNP^QiI^ for 24 hours, followed by outgrowth in non-selective YES (UT) for indicated times (hours) followed by either RNA-seq or testing for growth in PRD^DMI^ media. (B) WT cells were grown in YES (UT), PRD^DMI^ (control), or for 24 hours with FNP^QiI^ followed by washout and non-selective growth for indicated time (hours) prior to growth in PRD^DMI^ as shown in (A). Growth curves: average three replicates. Shading: variation. (C-G) Volcano plots from RNA-seq following 24h FNP^QiI^ treatment plus 0h (C), 6h (D), 12h (E), 24h (F) or 48h (G) recovery in YES post-treatment, samples compared to untreated cells. Genes with p-value <0.01 and fold change value ≥1.5 are indicated (dark grey). Representative DEGs are indicated in orange (up) and purple (down). (H) Heatmap depicting oxidative stress gene expression from RNA-seq in samples collected as in (A). (I) Venn diagram representing overlap between genes induced in FNP^QiI^ treatment and COSG. (J) (G) Growth curves: WT cells in YES alone (UT), or with addition of MND, PRD^DMI^ or MND+ PRD^DMI^. (K) As in (J) but with FLC^DMI^ instead of PRD^DMI^. MND: menadione. (L) Efflux of Rhodamine 6G (R6G) in WT or *qcr7Δ* cells untreated (UT, black) or treated with FNP^QiI^ (blue).

To investigate transcriptional changes induced by FNP^QiI^ we performed RNA-seq on cells 0, 6, 12, 24 and 48 hours after FNP^QiI^ washout (**Fig 4A**). Exposure to FNP^QiI^ for 24 hours prior to harvesting (0 hr) revealed a reversible downregulation of genes involved in oxidative phosphorylation (**Fig 4C-4G**). This group of repressed genes overlaps significantly with the previously reported mitonuclear retrograde (MNR) response, a signalling pathway coordinating the expression of mitochondrial and nuclear encoded mitochondrial proteins (S2A and S2B Fig) [47, 48]. FNP^QiI^ exposure activates MNR, decreasing expression of many nuclear genes encoding mitochondrial ETC components including Nde1, Qcr1, 2, 6-10, Cox4-6, 9, 12, 13, and Atp1-5, 7, 14, 16-20 (S2A and S2B Fig). In addition, genes that were upregulated upon FNP^QiI^ treatment also overlapped significantly with previously reported MNR genes (S2A and S2B Fig) [48]. Although the *S. pombe* MNR pathway is not well characterized, compared to *S. cerevisiae*, its activation likely limits ROS damage by dampening mitochondrial ETC activity [47, 48]. FNP^QiI^ also triggered expression of oxidative stress responding genes, which remained elevated 6 hours after FNP^QiI^ removal, thereafter, returning to normal (**Fig 4C-4H**). Specific analyses of transcripts by RT-qPCR confirmed the upregulation of the known oxidative stress response genes *bfr1*^+^, *pmd1*^+^ (encode ABC multidrug transporters) and *obr1*^+^ (encodes quinone oxioreductase) following FNP^QiI^ exposure and their return to normal by 6 hours (*pmd1^+^*) or 24 hours (*bfr1*^+^ and *obr1^+^*) (S2C Fig). Consistent with MNR activation by FNP^QiI^ treatment, a reciprocal transient decrease in expression of the MNR gene *nde1^+^*, which encodes mitochondrial NADH dehydrogenase, was also detected with recovery of full expression 24 hours after FNP^QiI^ removal (S2C Fig). *S. pombe* oxidative stress response genes have previously been shown to be induced by the ROS-generator menadione (MND) [49]. As with FNP^QiI^, exposure of wild-type cells to MND also suppressed PRD^DMI^ and FLC^DMI^-mediated growth inhibition (**Fig 4J** and **4K**). The finding that ETC-complex III inhibitors or a ROS generator promote azole^DMI^ resistance reinforces the concept that suppression of azole^DMI^ activity in *S. pombe* can be driven by oxidative stress induced by QoI/QiI antifungals.

Oxidative stress upregulates genes encoding detoxification activities and transmembrane transporters, and increases efflux through such transporters is a known mechanism of azole^DMI^ resistance in many fungi [43, 44, 49]. Thus, increased cellular efflux may explain how FNP^QiI^ renders cells more azole^DMI^ resistant. We measured the rate of Rhodamine 6G dye efflux as an indicator of overall efflux activity. When ETC-complex III function was impaired by either addition of FNP^QiI^ to wild-type cells or in *qcr7*Δ mutant cells, the Rhodamine 6G efflux rate was elevated (**Fig 4L**) [24]. We conclude that ETC-complex III-inhibiting fungicides such as QiIs trigger the oxidative stress response, thereby stimulating detoxification and efflux pump activities which decreases intracellular azole^DMI^ levels and consequently confers resistance to azole^DMI^ antifungals.

The trajectory of FNP^QiI^ -triggered transcriptional changes correlate with declining PRD^DMI^ resistance following FNP^QiI^ removal (**Fig 4A-4I**). Our finding that FNP^QiI^ exposure can enhance PRD^DMI^ resistance in fission yeast, an innocuous non-pathogenic model fungal system, raises the possibility that ETC-complex III inhibitor exposure in the environment could have collateral effects on non-target fungi with pathogenic potential.

### Exposure of the human fungal pathogen *Cryptococcus neoformans* to FNP^QiI^ enhances fluconazole resistance

*Cryptococcus neoformans* is an opportunistic basiodiomycete fungal pathogen and distantly related to the ascomycete *S. pombe* and is classified by the WHO as a critical priority pathogen [1, 2]. *C. neoformans* is found in soil, and associated with plant material and bird/pigeon droppings [18, 20]. Cryptococcal infections typically arise in immunocompromised individuals through airborne spore inhalation [50]. Fluconazole (FLC^DMI^) remains the only orally delivered and primary treatment for cryptococcosis, especially in resource-poor regions where advised combination therapy is less accessible [9, 10]. To assess whether prior exposure of *C. neoformans* to FNP^QiI^ affects azole^DMI^ sensitivity, we grew KN99 reference strain cells in a sublethal concentration of FNP^QiI^ for 24 hours prior to plating in the presence or absence of FLC^DMI^. FNP^QiI^ pretreatment alone did not affect cell viability but it induced a ten-fold increase in the number of FLC^DMI^ resistant colonies (**Fig 5A** and **5B**). As with *S. pombe*, FNP^QiI^ induced resistance of *C. neoformans* was transient as the frequency of FLC^DMI^-resistant colonies declined with longer outgrowth following FNP^QiI^ washout (**Fig 5B**). Relative FLC^DMI^ resistance following FNP^QiI^ washout was also assessed by continual measurement of optical density in multiple microwell cultures [51], both *C. neoformans* KN99 and clinical isolate IFN80 (VNBII lineage [52]) showed increased FLC^DMI^ resistance that again declined with time after FNP^QiI^ removal (**Fig 5C** and **5D**). These observations suggest that residual FNP^QiI^ present in field soil, pigeon droppings, or plant material has the potential to increase *C. neoformans* resistance to azole-based antifungals in resident fungi while it remains active.

**Fig. 5.**
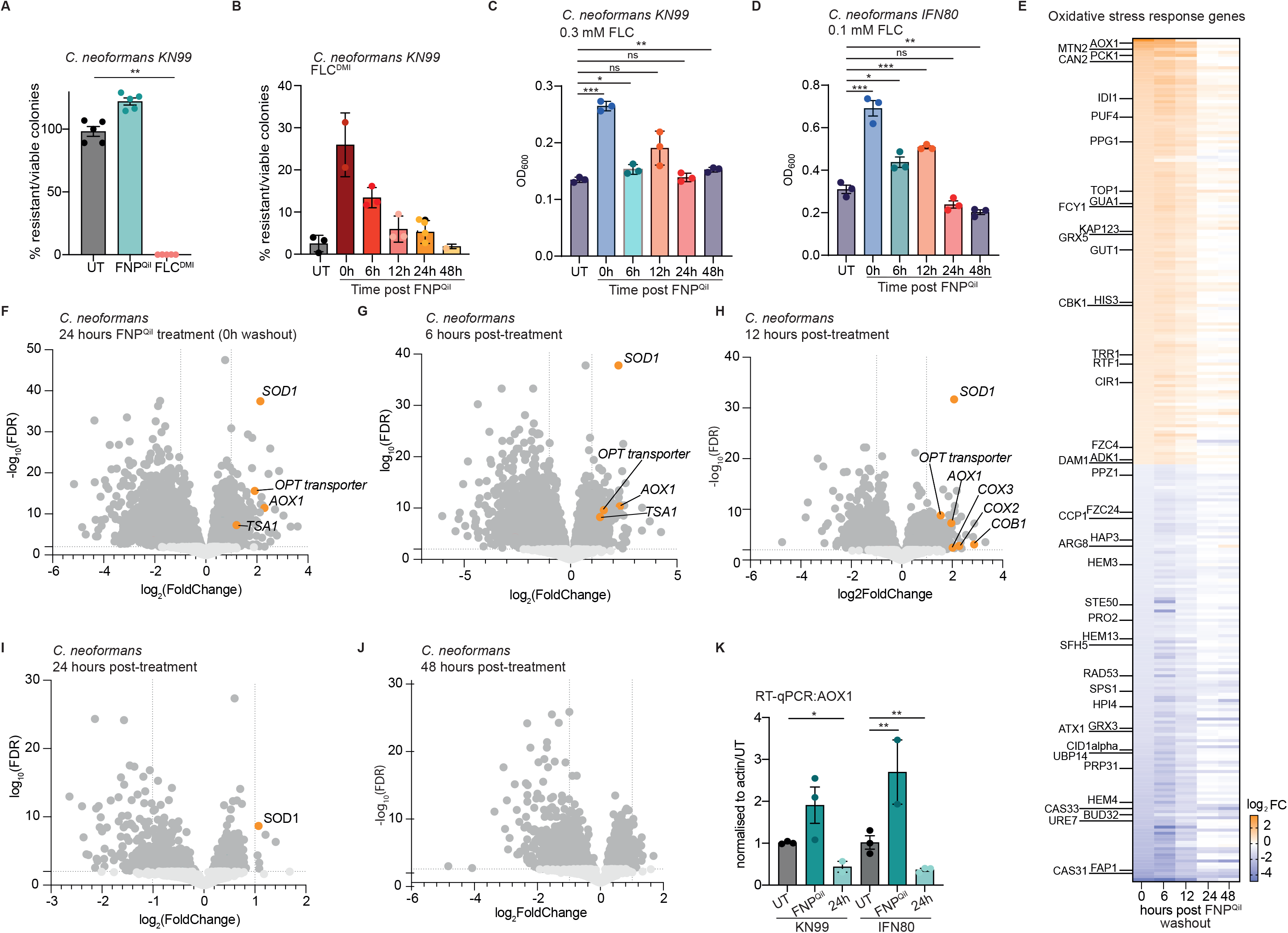
FNP^QiI^ treatment suppresses FLC^DMI^ sensitivity of *C.neoformans*. (A) Colonies formed by KN99 cells on YPD alone (UT) or with addition of FLC^DMI^ or FNP^QiI^ expressed as a percentage of viable colonies grown on YPD alone. (B) Percentage of viable colonies formed by KN99 cells on FLC, comparing untreated (UT) cells and those grown for 24 hours in FNP^QiI^ followed by washout and non-selective growth for indicated time (hours) (as in Fig 4A). (C) Microtitre liquid growth assay using KN99 showing OD_600_ comparing untreated (UT) cells and those grown for 24 hours in FNP^QiI^ followed by washout and non-selective growth for indicated time (hours) (as in Fig 4A). (D) As (C) using the IFN80 clinical isolate. (E) Heatmap depicting annotated oxidative stress gene regulation in samples collected at 0, 6, 12, 24 and 48h (as B, F-J). (F-J) Volcano plots from RNA-seq following 24h FNP^QiI^ treatment plus 0h (F), 6h (G) or 12h (H), 24h (I) or 48 h (J) YPDA recovery after washout, samples compared to untreated KN99 cells. Genes with p-value <0.01 and fold change value ≥1.5 are indicated (dark grey). Representative DEGs are indicated in orange (up). (L) RT-qPCR analysis of *AOX1* expression in *C. neoformans* KN99 and clinical isolate IFN80 untreated cells (UT) compared to cells exposed to FNP^QiI^ for 24 hours or cells given 24 hours recovery in non-selective YPDA following FNP^QiI^ washout. Significance: *Student’s t-test; **p<0.01, *p<0.05*.

To determine how FNP^QiI^ exposure might increase *C. neoformans* resistance to FLC^DMI^, we performed RNA-seq on KN99 cells grown with or without FNP^QiI^ for 24 hours and, following washout, grown in its absence for 6, 12, 24 and 48 hours (as in **Fig 4A**). Alternative oxidase (*AOX1*) gene expression was highly induced by FNP^QiI^ treatment and remained high 6-12 hours after FNP^QiI^ withdrawal (**Fig. 5E-J**). Both *S. cerevisiae* and *S. pombe* lack genes encoding alternative oxidases, but *AOX1* expression is known to be induced by oxidative stress in *C. neoformans*, and *Candida albicans* where its deletion leads to increased FLC^DMI^ sensitivity [53–55]. Alternative oxidases are known to bypass ETC-Complex III, potentially by decreasing ROS generation thereby providing protection in *C. neoformans* and other fungi [54, 55]. The *SOD1* (Superoxide dismutase) and *TSA1* (Thiol-specific antioxidant protein) genes, that are induced by oxidative stress in *C. neoformans* [56], also displayed increased expression in response to FNP^QiI^, with expression decreasing to untreated levels after 12 hours (**Fig 5E-5J**) consistent with FNP^QiI^ causing oxidative stress via mitochondrial dysfunction.

Unexpectedly, the expression of genes encoding efflux pumps was not significantly altered indicating that the FNP^QiI^-mediated fluconazole resistance mechanism deployed by *C. neoformans* differs from the Pap1-efflux axis activated by FNP^QiI^ in *S. pombe*. Indeed, although fungal stress response pathways are generally conserved, interconnections are wired differently in distinct species perhaps reflecting their differing niches and related but different responses [57].

We also found *AOX1* expression to be induced by FNP^QiI^ in the less laboratory adapted *C. neoformans* clinical isolate IFN80 (**Fig 5L**). Thus, pretreatment of *C. neoformans* with FNP^QiI^ induces a limited stress response, where Aox-adapted metabolism and associated alteration in cell wall composition, uptake channels and detoxification activities may restrict intracellular fluconazole levels [53, 57]. Attenuated mitochondrial functions were recently found to increase FLC resistance in *C. neoformans*. It is possible that the resulting increase in acetyl-COA availability, the first step in ergosterol biosynthesis, upon ETC blockage may simply compensate for Erg11 inhibition by FLC [27, 58]. Irrespective of the exact mechanism, environmental exposure of *C. neoformans* to QiI fungicides likely creates transient conditions that facilitate the emergence of azole resistance, thereby posing an increased risk to humans exposed to these more resistant fungal populations.

### FNP^QiI^ treatment increases resistance of *Zymoseptoria tritici* to azole^DMI^-based antifungals

QiI and QoI fungicides are used to limit fungal-borne crop diseases and are commonly used in combination with azole^DMI^-based Erg11/Cyp51 inhibitors to tackle Septoria Tritici/Leaf Blotch caused by the ascomycete fungus *Z. tritici* which damages foliage and reduces grain yields [12, 14, 16]. *Z. tritici* overwinters on wheat stubble from the preceding season, providing a source of wind-borne spores that infect newly developing plants [59]. Currently, combinations of QiIs, QoIs, SDHIs and azoles^DMI^ are used at different stages of plant development to hinder fungal infections, with a mix of FNP^QiI^ and PRD^DMI^ sprayed on wheat fields annually upon flag leaf emergence [12]. The standard *Z. tritici* reference strain IPO323 fails to grow in PRD^DMI^ whereas pre-treatment with a sub-lethal concentration of FNP^QiI^ reduced viability by half (**Fig 6A** and S3A Fig). In contrast, *Z. tritici* growth was greater on plates containing both FNP^QiI^ and PRD^DMI^ than on those containing PRD^DMI^ alone (**Fig 6B**). Low residual FNP^QiI^ remaining in soil or plant debris could potentially influence *Z. tritici* resistance to azole^DMI^ treatment. To mimic such conditions, we exposed *Z. tritici* IPO323 cells to FNP^QiI^ for 24 hours prior to plating in the presence or absence of PRD^DMI^. This pre-treatment resulted in a three-fold increase in the number of PRD^DMI^ colonies formed (T0 days). As with *S. pombe* and *C. neoformans*, FNP^QiI^-stimulated *Z. tritici* resistance to PRD^DMI^ was transient and returned to control levels after 3 days outgrowth in the absence of FNP^QiI^ (**Fig 6C**).

**Figure 6.**
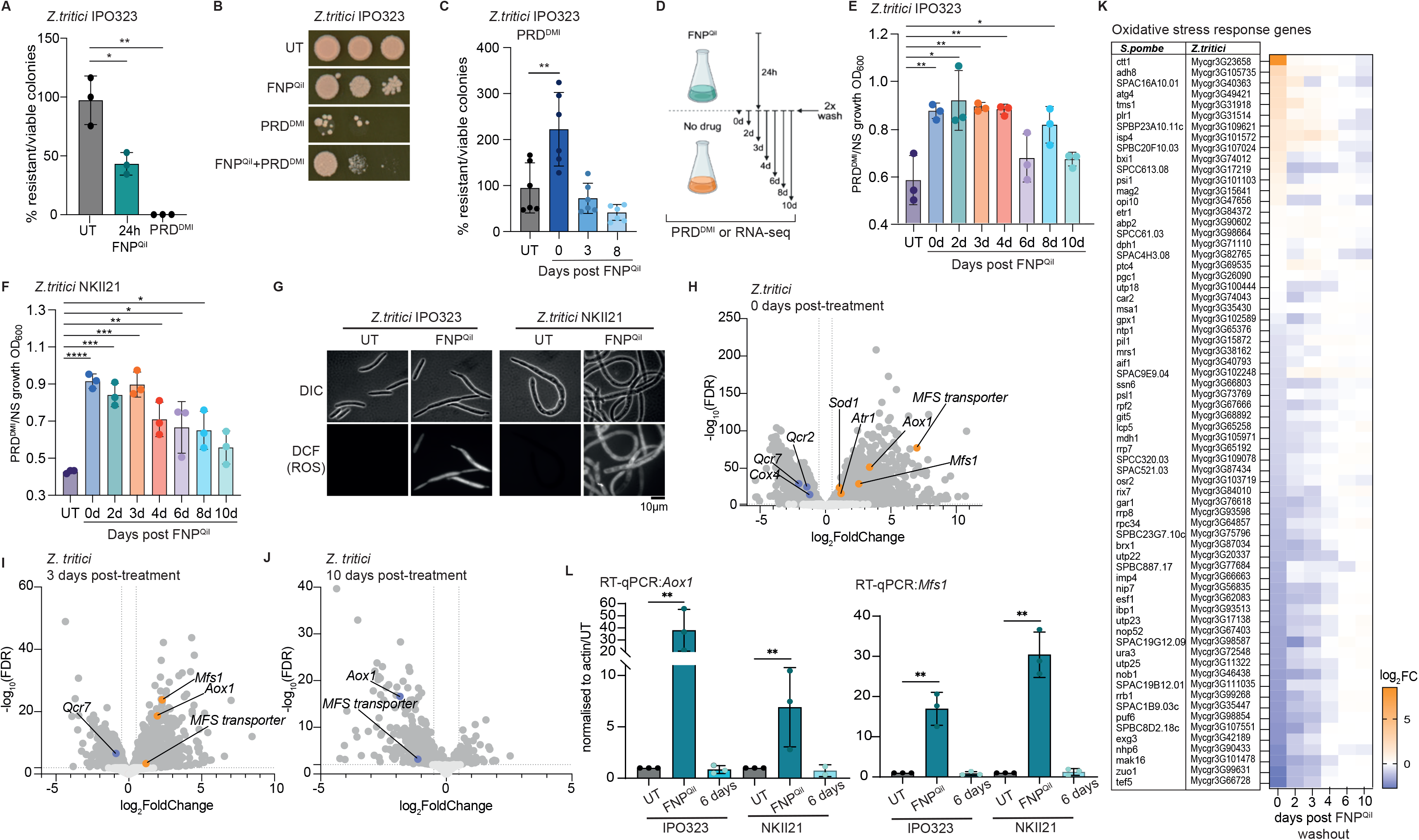
FNP^QiI^ fungicide exposure increases azole^DMI^ resistance in *Z. tritici*. (A) Viability of *Z. tritici* IPO323 grown untreated (UT) or in FNP^QiI^ for 24 hours prior to plating on non-selective YPD, or UT cells plated directly on PRD^DMI^. (B) Comparative growth assays on YPD alone, or with addition of PRD^DMI^, FNP^QiI^ or PRD^DMI^+FNP^QiI^ (C) Percentage survival on PRD^DM^ after 24 hours FNP^QiI^ exposure followed by 0, 3, or 8 days recovery in YPD. (D) Schematic of experimental set-up. IPO323 cells were pre-exposed to FNP^QiI^ for 24 hours, then washed and grown in non-selective YPD for indicated days followed by RNA-seq or growth in PRD^DMI^. (E-F) Growth of IPO323 (E) or NKII21 (F) cells treated as in (D). (G) DCF ROS detection. WT IPO323 or NKII21 cells untreated (UT) or treated with FNP^QiI^ incubated in DCFH-DA ROS-indicator. (H-J) Volcano plots from RNA-seq following 24h FNP^QiI^ treatment plus 0h (H), 6h (I) or 24h (J) YPD recovery following washout, samples compared to untreated IPO323 cells. Genes with p-value <0.01 and fold change value ≥1.5 are indicated (dark grey). Representative DEGs are indicated in orange (up) and purple (down). (K) Heatmap depicting annotated oxidative stress gene regulation over 0-10 days recovery (D). *Z. tritici* and equivalent *S. pombe* accession codes are shown. (L) RT-qPCR comparing *Aox1* or *Mfs1* expression in untreated IPO323 or NKII21 cells (UT) to cells exposed to FNP^QiI^ for 24 hours or following 6 days’ YPD recovery. Significance: *Student’s t-test **<0.01*.

*Z. tritici* blastospores are irregularly shaped and can form pseudo-hyphae under laboratory conditions. Comparison of colony forming unit counts on plates with liquid optical density measurements from the same cultures for both *Z. tritici* IPO323 and Hungarian field isolate NKII21 [51, 60] showed a linear relationship, validating microwell culture optical density readings for measuring relative resistance (S3B and S3C Fig). Following FNP^QiI^ exposure for 24 hours and subsequent wash-out, *Z. tritici* cells were incubated non-selectively for 0-10 days and their ability to grow in PRD^DMI^ assessed at each timepoint (**Fig 6D-6F**). FNP^QiI^ exposure consistently enhanced *Z. tritici* survival in PRD^DMI^ with resistance declining following longer periods of recovery after FNP^QiI^ treatment (**Fig 6E and 6F**).

Treatment of both IPO323 and NKII21 *Z. tritici* cells with FNP^QiI^ for 24 hours resulted in ROS accumulation (**Fig 6G**). Moreover, RNA-seq analyses revealed that this exposure of IPO323 cells to FNP^QiI^ caused downregulated expression of genes involved in oxidative phosphorylation, similar to the *S. pombe* mitonuclear retrograde response (MNR) (**Fig 6H-6J**). FNP^QiI^ also induced a group of *Z. tritici* genes encoding orthologs of proteins known to be induced by oxidative stress in *S. pombe* (**Fig 6K**), as well as MFS multidrug transporters such as Mfs1 (**Fig 4H and 4I**). Furthermore, as in *C. neoformans,* the gene encoding Aox1 was also induced (**Fig 4H and 4I**). Expression of several oxidative stress response genes remained elevated for 4 days after FNP^QiI^ removal from *Z. tritici* IPO323 cells and the period of their increased expression coincided with increased PRD^DMI^ resistance (**Fig 6E, 6H-6K and S3D-S3F Fig**). Transient induction of *Aox1* and *Mfs1* in response to FNP^QiI^ was confirmed by RT-qPCR for both IPO323 and NKII21 (**Fig. 6L**). Thus, exposure of *Z. tritici* to FNP^QiI^ induces oxidative stress and stress-response genes which temporarily enhances its resistance to recommended azole-based DMIs such as prothioconizole-desthio that are often also prevalent in fields due to crop spraying.

## Discussion

We have shown that sublethal exposure to the fungicide FNP^QiI^ suppresses the effectiveness of azole^DMI^ antifungals in the model fungus *S. pombe* and in the fungal pathogens *C. neoformans and Z. tritici*. In *S. pombe*, suppression arises from oxidative stress generated by FNP^QiI^ inhibition of the ETC, which activates the Pap1-dependent oxidative stress transcriptional program, including the upregulation of efflux pumps capable of lowering intracellular azole accumulation (**Fig. 7**). A related oxidative stress resistance mechanism presumably mediated by ZtYap1 (Pap1 orthologue) appears to operate in *Z. tritici* [61], which may also respond to the observed FNP^QiI^-induced ROS to mediate azole^DMI^ resistance.

**Figure 7.**
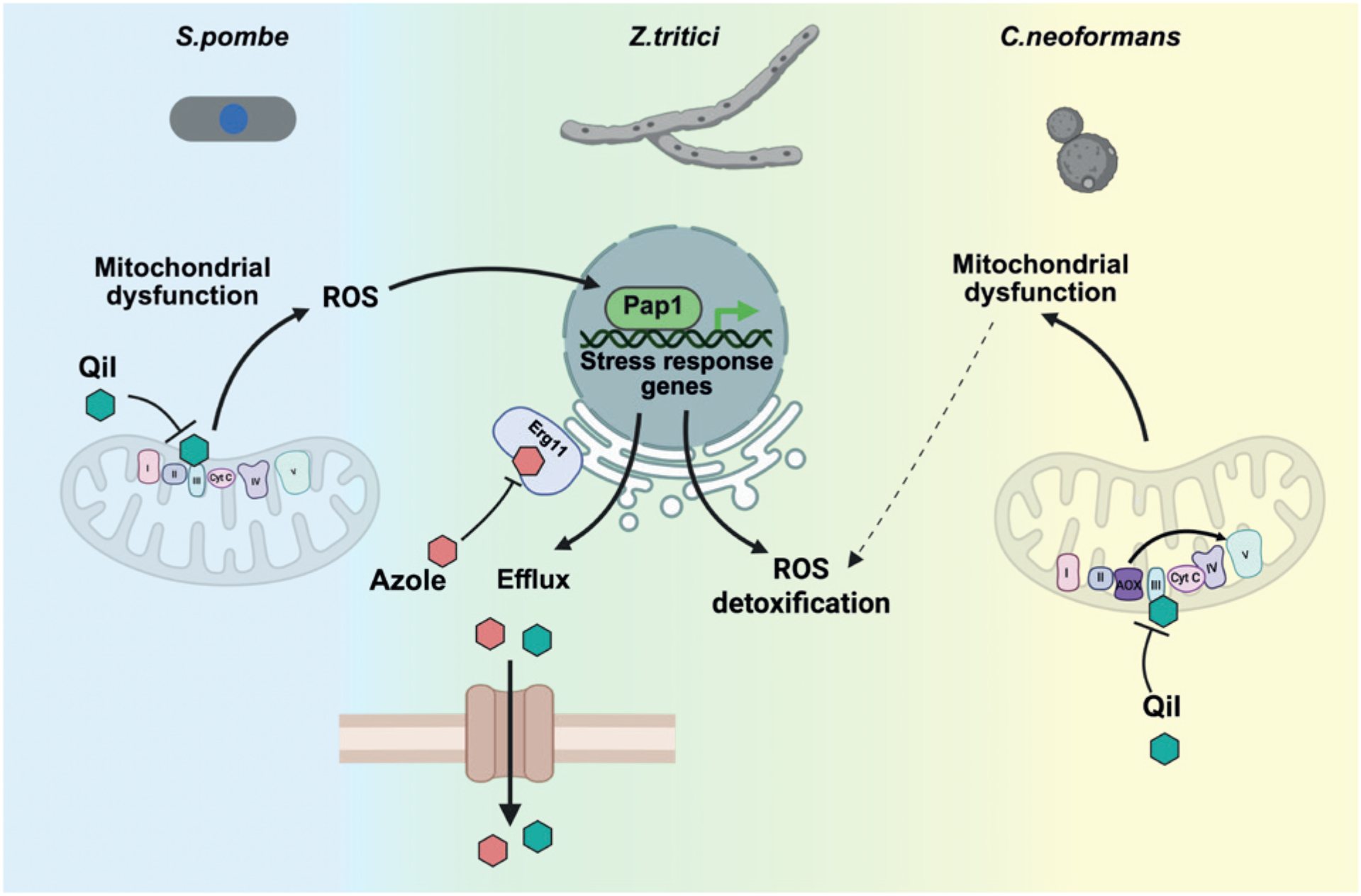
Model of stress responses contributing to fungal resistance mechanisms. Qil targets mitochondrial ETC complex III in *S. pombe* and *Z. tritici* causing mitochondrial dysfunction and the generation of reactive oxygen species (ROS). In *S. pombe* elevated ROS activates the transcription factor Pap1, which induces the expression of stress response genes involved in ROS detoxification and drug efflux. These responses limit intracellular accumulation of azoles and cells develop increased resistance to azoles. A subset of oxidative stress genes that are regulated in *S. pombe* are also observed in *Z. tritici*. *Z. tritici* and *C. neoformans* also contain an additional enzyme, alternative oxidase (AOX), which allows electrons to bypass the main cytochrome pathway to mitigate the generation of ROS. *Z. tritici* utilises both efflux and AOX to allow cells to develop azole resistance, however *C. neoformans* appears to achieve resistance primarily via AOX and detoxifying enzymes. In both cases Qil impairs the mitochondrial respiratory chain. This model highlights a conserved mechanism of Qil action across diverse fungi and underscores the protective role of Pap1-regulated stress responses and AOX in mediating fungal resilience to antifungal agents

In contrast, *C. neoformans* mounts a distinct response upon FNP^QiI^ exposure. In common with many plants and other fungi, both *C. neoformans* and *Z. tritici* possess alternative oxidases, which can bypass ETC-Complexes III/IV to maintain respiration; ATP production is reduced but ROS accumulation is limited [53, 54]. Following FNP^QiI^ treatment, Aox is strongly induced in both species, presumably acting to mitigate oxidative stress, enhancing resistance and cell survival. In *Z. tritici*, Aox-mediated protection is accompanied by upregulation of efflux pumps, further counteracting oxidative stress and consequently conferring azole^DMI^ resistance. Unexpectedly, efflux pump expression was largely unchanged by FNP^QiI^ treatment of *C. neoformans*, suggesting that azole^DMI^ resistance arises primarily through detoxification and the Aox-driven metabolic adaptations that accompany it, which themselves reduce ETC-coupled ROS generation [53].

These findings highlight the potential for agricultural fungicide use to have consequences that reach beyond the selection of resistant crop fungal pathogens, and indicate that certain antifungal combinations may be counterproducitve. Antifungal compounds differ in their half-lives and may persist following application to growing crops and remain in soil, previous season’s stubble, or field margins, exposing resident fungal populations to residual concentrations of fungicides, such as QiIs and QoIs. Such exposure may prime transient azole^DMI^ resistance, allowing fungal cells which would otherwise be eliminated by azole^DMI^ treatment to survive and thereby increasing the opportunity for resistance conferring mutations to arise and become established in populations. More broadly, the widespread application of agricultural fungicides may contribute to the depletion of beneficial fungi, with detrimental effects on soil quality, the selection of fungicide resistant crop pathogens and the conditioning of resistance in potential opportunistic human fungal pathogens [I did suggest a Nat Comm ref for this if wanted].

Resistance conditioning is of particular concern because human opportunistic pathogens, including *C.neoformans* and *A.fumigatus,* are associated with plant material and may therefore encounter fungicide residues in the environment. The suppression of azole antifungal effectiveness by pretreatment or in combination with ETC-complex III inhibitors, as demonstrated here, raises the possibility that such environmental exposure could enhance the survival of these fungi during subsequent azole^DMI^ treatment, facilitating the emergence and selection of clinically relevant antifungal resistance.

Moreover, the overlap between compounds used in both clinical and agricultural settings no longer applies to just azole-based antifungals. The agricultural dihyroorotate dehydrogenase (DHODH) inhibitors Ipflufenoquin and Quinofumelin are directed to the same target as the drug Olorofim, developed to treat human aspergillosis [62]. This raises the possibility that exposure to agricultaral DHODH inhibitors could select for resistance before the drug is employed to combat human disease. DHODH itself is a key mitochondrial enzyme, so its inhibition might perturb mitochondrial redox balance and thereby, as with ETC-complex III inhibitors, suppress the activity of other classes of antifungal drugs.

Although we have identified clear antagonism between two antifungals used to control a major wheat fungal pathogen, exposure of crops to diverse agrochemical pesticides, including those that impact insect development, raises the possibility of further suppressive interactions [63]. The widespread use of fungicides in agriculture is essential to protect crop yields and maintain food security; however, these benefits must be balanced against potential consequences for environmental and human health. In particular the persistence of ETC-complex III inhibitors in treated environments and their potential to condition antifungal resistance highlight the need for careful management of their agricultural use. Such management should seek to preserve the effectiveness of crop protection strategies whilst minimising the selection and spread of antifungal resistance in environmental fungi, including those with the potential to cause human disease.

## Acknowledgements

The authors thank Allshire lab members and Donal O’Carroll for comments and discussion. We acknowledge the Discovery Research Platform for Hidden Cell Biology for assistance provided by David Kelly and Toni McHugh through the Light Microscopy Core and by Shaun Webb through the Bioinformatics Core.

## Author contributions

Conceptualization: RCA, ALP, SAW

Methodology: ALP, PT, SMP, SAW

Investigation: ALP, SMP, RY, SAW

Visualization: RCA, ALP, SMP, SAW,

Funding acquisition: RCA, ALP

Project administration: RCA, ALP, SAW

Supervision: RCA, ALP, SAW

Writing – original draft: RCA, ALP, SAW, SMP,

Writing – review & editing: RCA, ALP, SAW, SMP

## Data and materials availability

All data are available in the main text or the supplementary materials apart from the primary RNA-seq data. RNA-Seq data is available to reviewers at GEO using the code: Original uncropped microscopy and plate images are available on request. Excel files of growth curve, flow cytometry, efflux measurements, qRT-PCR data are available on request.

## Materials and Methods

### *S. pombe*, *C. neoformans*, *Z. tritici* strains growth and manipulation

Strains used in this study are listed in S1 Table. Mutation of *erg11* was performed using SpEdit CRISPR-Cas9 methodology ^52^ using oligonucleotides listed in S2 Table. Chemicals and reagents were obtained from Merck unless indicated otherwise. Azole antifungal drugs were obtained from Ehrenstorfer. Wild-type *Schizosaccharomyces pombe* ED972 cells were grown at 32°C in Yeast Extract (Difco) + elements (YES) medium: 0.5% w/v yeast extract, 0.2 g/l adenine, 0.2 g/l arginine, 0.2 g/l lysine, 0.2 g/l histidine, 0.2 g/l uracil, 0.2 g/l leucine (doubling time 2 hours 10 minutes). For YEG media, the glucose in YES media was replaced with 3% glycerol + 0.1% glucose. Wild-type *Zymoseptoria tritici* IPO323 was grown at 18°C in YPD (doubling time 8 hours), the *Z. tritici* Hungarian 2008 field isolate NKII21/CBS 123916 was obtained from the Westerdijk Fungal Biodiversity Institute, Utrecht, The Netherlands; https://wi.knaw.nl/fungal_table. *Cryptococcus neoformans* KN99 was grown at 30°C in YPDA (doubling time 1 hour 52 minutes), the recent clinical isolate IFN80 was obtained from Tihana Bicanic, St George’s University of London. Concentrations of agents used were as follows. *S.pombe*: 135ug/ml fluconazole (FLC^DMI^); 0.4 μg/ml Prothioconazole-desthio (PRD^DMI^); 0.01 mg/ml fenpicoxamide-phenol (FNP^QiI^); 5 μg/ml mefentrifluconazole (MFZ^DMI^); 0.25 μg/ml antimycin A (ANT); 0.02 mg/ml azoxystrobin (AZS^QoI^), 12.5μg/ml thiabendazole (TBZ), 0.5M potassium chloride (KCl). *C.neoformans*: 45.9ug/ml fluconazole (FLC^DMI^); 0.05 mg/ml fenpicoxamide-phenol (FNP^QiI^). *Z. tritici*: 4 ng/ml Prothioconazole-desthio (PRD^DMI^); 1 μg/ml fenpicoxamide-phenol (FNP^QiI^).

### Growth curves

Cells were grown in Tecan Sunrise microwell plate readers using the following parameters. *S. pombe* (4×10^4^ cells/well) was grown in 100 μl of media in a clear flat bottom 96-well plates (Costar) at 32±2°C with continuous shaking. Prior to measurement, plates were allowed to settle for 5 seconds. Data was collected using Magellan software. Measurement was conducted for 48 hours with OD_600_ measurements every ∼15 minutes. All growth curves were performed at least in triplicate and analysed in Excel, then visualized in GraphPad Prism.

### Serial dilution comparative growth assays

Equal numbers of cells were serially diluted four-fold in sterile water and spotted on indicated agar plates using a pinning device. Plates were incubated at appropriate temperatures (*S.pombe;* 32°C, *Z.tritici;* 18°C, *C.neoformans;* 30°C) for 3 to 15 days before imaging with an Epson flat-bed scanner at 600 dpi.

### TPF detection assay for respiratory competence

The 2,3,5-triphenyltetrazolium chloride (TTC) dye is normally reduced to red 2,3,5-triphenyltetrazolium formazan (TPF) by respiring cells. TPF detection assays were performed as previously ^20^. Plates were grown for 3 days and overlayed with 1% UltraPure™ Low Melting Point Agarose containing 0.5 mg/ml TTC in 100 mM sodium phosphate. Cell respiration was evaluated by colony color after 24 h incubation with red colony color indicative of functional respiration. If respiration is defective, the color is unchanged.

### Fluorescence microscopy

#### GFP-Pap1 detection and measurement

Cells were grown to log phase (5 X10^6^/ml) and fixed for 10 minutes at room temperature with 3.7% formaldehyde (Merck, F8775). Cells were washed twice in PEM (100 mM PIPES pH7, 1 mM MgCl_2_, 1 mM EDTA), then resuspended in PEMS (PEM plus 1.2M sorbitol). After pelleting cells were incubated for 5 minutes at room temperature in PEMS containing 1% Triton-TX100 (Merck, X). Samples were washed once in PEMS, followed by one wash in PEM and incubated in PEM/0.1% sodium azide and DAPI for 5 minutes. Cells were then washed in PEM/0.1% sodium azide and analysed by imaging.

Cells were mounted in VECTASHIELD medium (Vector Labs). Widefield fluorescence images were taken with a Nikon Ti2 microscope equipped with a Prime 95B camera (Teledyne Photometrics, Tucson, AZ 85706), using an Apo TIRF 100x / 1.49 oil objective; DAPI filter, excitation 377/26, emission 432/36; GFP filter, excitation 474/26 emission 525/40 and Spectra X light-source (Lumencor, Beaverton, OR 97006 USA). Analysis was performed using FIJI. The nuclear area was determined by thresholding the DAPI channel (MaxEntropy method). The GFP signal in each cell was measured both in the nucleus and cytoplasm (nuclear region of interest (ROI) manually moved into cytoplasm). To determine Pap1 nuclear localisation this formula was employed (nuclear - cytoplasmic GFP signal)/(nuclear + cytoplasm GFP signal). At least 200 cells were analysed per sample. Significance of the difference between samples was determined using Student’s t-test.

### ROS detection with DCFH-HA sensor

The ROS sensor dichlorodihydrofluorescein diacetate (DCFH-DA) is converted to fluorescent 2′,7′-dichlorofluorescein (DCF) by intracellular ROS. Cells were stained for the presence of ROS using a method adapted from ^20^. Briefly, log-phase cells (7 x 10^7^) grown in YES were harvested by centrifugation, washed twice with PBS then incubated in PBS for 30 min at room temperature. Cells were incubated in 10 μM DCFH-DA for 30 min and analysed by imaging or flow cytometry.

For imaging of resulting fluorescent intracellular DCF, cells were identified using brightfield illumination to ensure no pre-exposure of cells to 488 nm light, which itself induces ROS, prior to imaging. Cells were imaged on a Zeiss Axio Imager A2 fluorescence microscope with a 100X, 1.4-NA Plan-Apochromat objective, Prior Scientific filter wheel, Chroma 86000 filter set controlled by and illumination by HBO100 mercury bulb. Images were acquired using a Photometrics Prime sCMOS camera (Teledyne Photometrics, Tucson, AZ, 85706) controlled using Micromanager software.

### ROS quantification by Flow Cytometry

Levels of DCFH-DA converted to fluorescent DCF by intracellular ROS was measured by flow cytometry with a Thermo-Fisher Attune NxT flow cytometer as detailed ^20^, 488nm laser excitation, 4 channels (channel 1 = 530/30, channel 2 =574/26, channel 3 = 695/40, channel 4 = 780/60). Analyses and graphs were produced with FLowJo Software (FlowJo, Ashland, OR, 97520, USA). 100,000 cells were analysed for each sample.

### Rhodamine 6G efflux assay

Efflux was measured using the following protocol adapted from ^20^. All incubations were carried out at room temperature (∼25°C). Cells were grown in YES and ∼2.5 x10^8^ log-phase cells were harvested by centrifugation, washed twice with 25 ml PBS and incubated in 25 ml PBS for 30 min to deplete glucose and prevent rapid efflux of Rhodamine 6G (R6G) during uptake/loading. Pelleted cells were resuspended in PBS containing 10 μM R6G and incubated for 30 minutes in the dark. Cells were washed twice in PBS, pelleted and then YES (glucose rich media) added to allow efflux to commence. Samples were taken at 2 minute intervals and centrifuged at 13,000 rpm for 45 seconds. The supernatant containing effluxed fluorescent R6G were loaded to a black 96-well plate (Costar) and R6G fluorescence was measured with a Spectramax M5 Multimode plate reader (Molecular Devices, Wokingham), using SoftMax Pro / SpecMaxMT software, and excitation at 525 nm, auto cut-off, emission 565 nm.

### RNA extraction and RT-qPCR

Log phase cultures were harvested and total RNA extraction was performed either using Monarch Total RNA Miniprep Kit (New England Biolabs T2010S) for *S.pombe,* or Direct-zol RNA Miniprep kit (Zymo Research R2050) for *Z.tritici* and *C.neoformans.* Extractions were performed according to the manufacturer’s instructions. For *S. pombe* contaminating DNA was removed by 1 hour incubation with Turbo DNAse (Invitrogen) followed by repeating washes and elution as per manufacturer’s instructions. Quantification of RNA was performed using NanoDrop (ThermoFisher Scientific) or 2100 Bioanalyzer (Agilent) depending on downstream application. To determine RNA transcript levels from genes of interest, quantitative reverse-transcriptase PCR (RT-qPCR) was performed. LunaScript RT SuperMix Kit (New England Biolabs E3010L) was used for complementary DNA (cDNA) synthesis from 400ng of RNA, following manufacturer’s protocol. Quantitative analysis by qPCR was carried out using a LightCycler (Roche) and LightCycler 480 SYBR Green Master Mix (Roche). Oligonucleotide sequences are detailed in Table S2.

### RNA-sequencing (RNA-Seq)

Log phase cultures were harvested from biological triplicates of indicated cells. Following total RNA extraction, mRNA selection was performed using NEB Next Poly(A) mRNA Magnetic Isolation Module (E7490) and cDNA libraries were prepared with NEBNext Ultra™ II Directional RNA Library Prep Kit for Illumina (New England Biolabs E7760) according to manufacturer’s protocol. Libraries were assessed for quality using an Bioanalyzer High Sensitivity DNA Assay Chip and Bioanalyzer instrument (Agilent Technologies). After pooling, libraries were sequenced by the Edinburgh Clinical Research Facility using the Illumina NextSeq platform. Raw reads were quality assessed with FastQC and aligned and mapped to the appropriate reference genome (*Schizosaccharomyces pombe* 972, ASM294v2; GCF_000002945.1; *Zymoseptoria_tritici* MG2, GCA_000219625.1; *Cryptococcus neoformans* var grubii h99, GCA_000149245, CNA3) using STAR RNA-seq aligner and SAMtools [64, 65]. Salmon was used to quantifying the expression of transcripts and differential expression analysis was carried out using DEseq2 [66]. Volcano plots and heat maps were generated using GraphPad Prism.

---

**S1 Fig.**
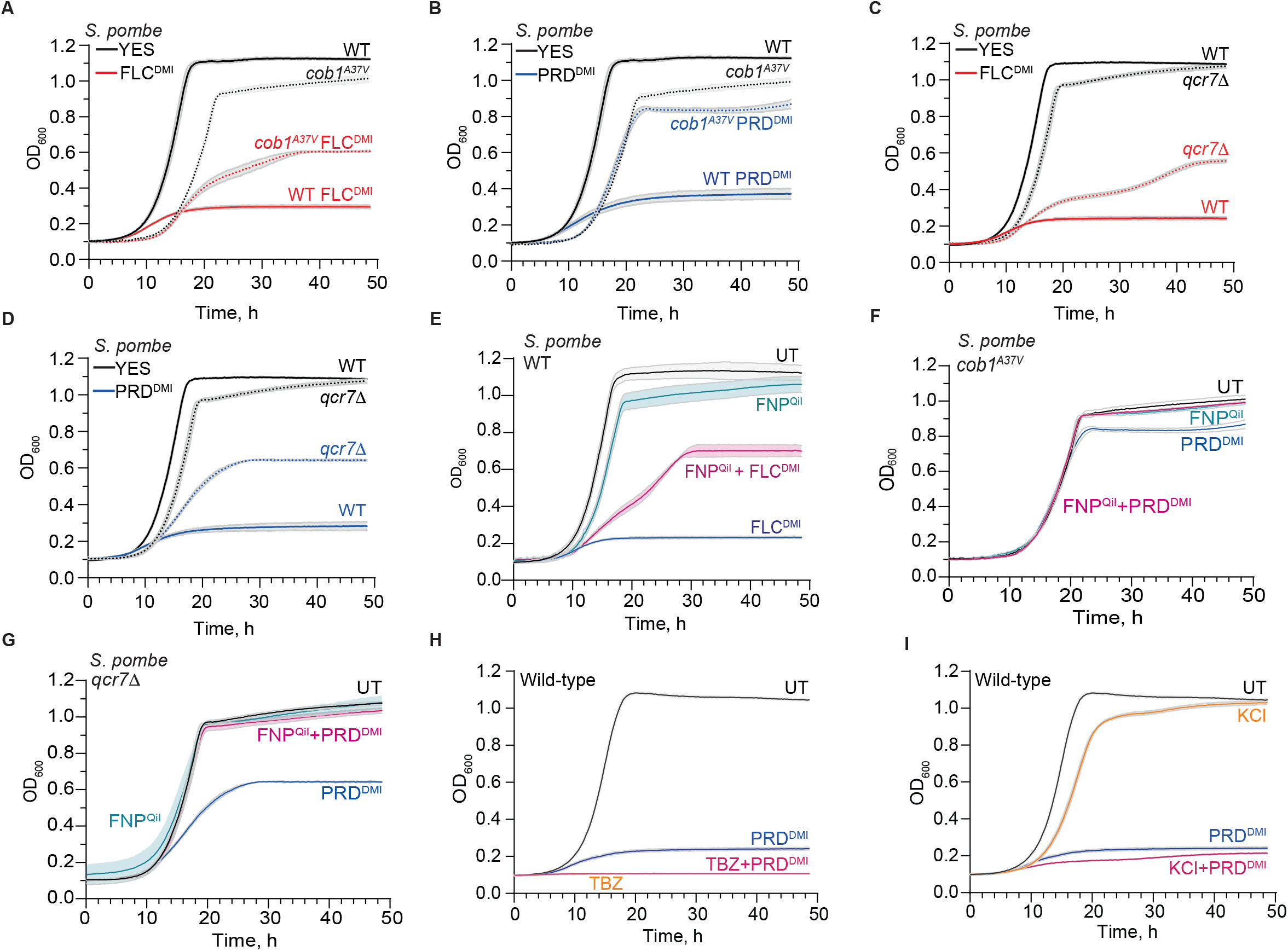
ETC-Complex III Qi and Qo inhibiting fungicides increase azole^DMI^ resistance of *S. pombe*. (A-D) Growth curves: WT cells in YES alone (UT, untreated), or with addition of FLC^DMI^ (red) or PRD^DMI^ (blue) compared to, c*ob1^A37V^* or *qcr7Δ*. (E) Growth curves: WT cells in YES alone (UT, black), or with addition of FNP^QiI^ (teal), FLC^DMI^ (blue) , or FNP^QiI^+FLC^DMI^ (dark pink). (F, G) Growth curves: *cob1^A37V^* (F) or *qcr7Δ* (G) cells in YES alone (UT), or with addition of FNP^QiI^, PRD^DMI^ or FNP^QiI^+PRD^DMI^. (H) Growth curves: WT cells in YES alone (UT, black), or with addition of PRD^DMI^ (blue), TBZ (orange) , or TBZ+PRD^DMI^ (dark pink). (I) Growth curves: WT cells in YES alone (UT, black), or with addition of PRD^DMI^ (blue), KCl (orange), or KCl+PRD^DMI^ (dark pink). Extended Data Figure 1. All panels: average three replicates. Shading: variation. FLC^DMI^: Fluconazole; PRD^DMI^: prothioconazole-desthio; FNP^QiI^: fenpicoxamid-phenol; TBZ: Thiabendazole; KCl: potassium chloride.

**S2 Fig.**
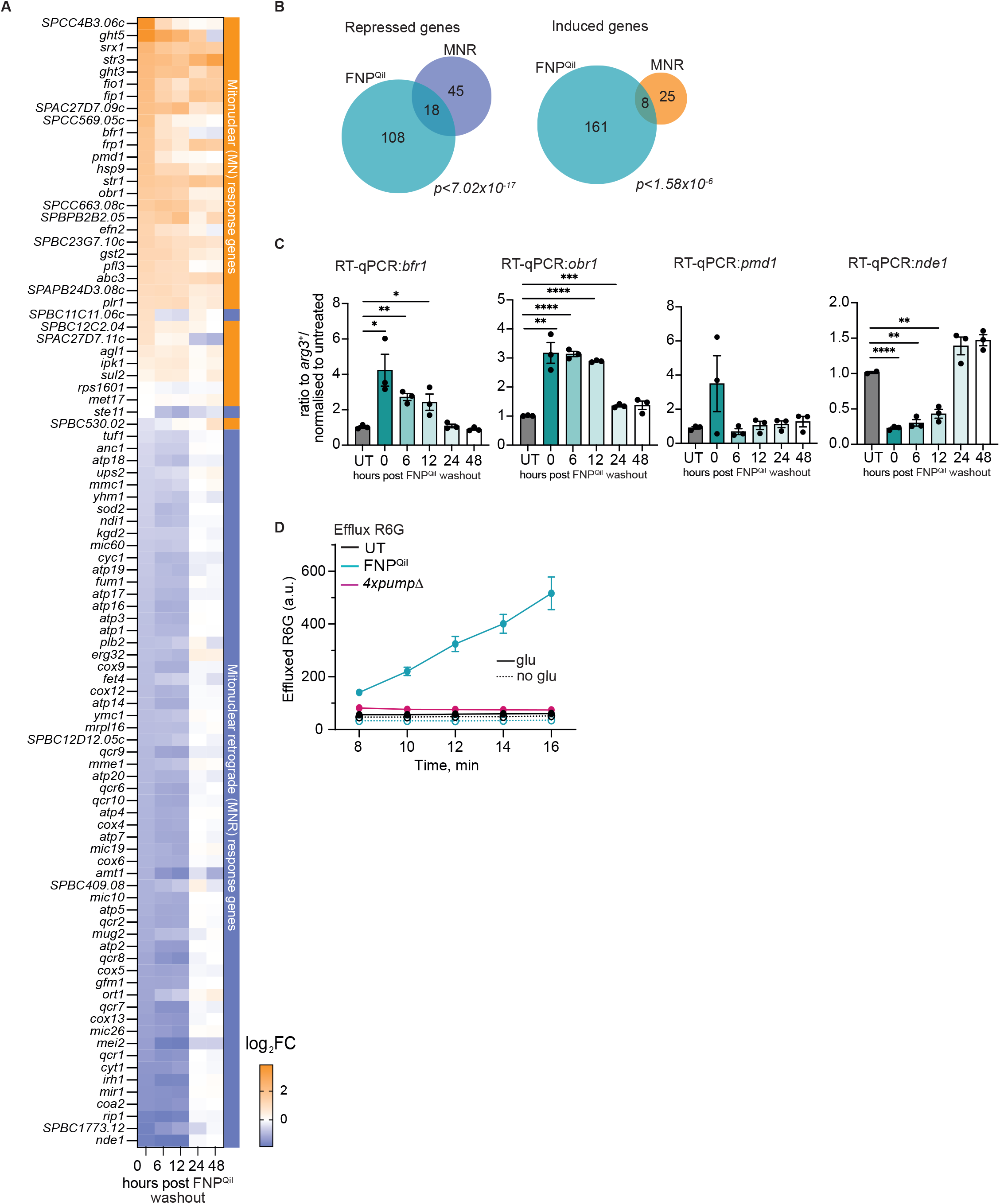
ETC-Complex III inhibitors induce mitochondrial dysfunction and activate the mitonuclear retrograde response. (A) Heatmap depicting mitonuclear retrograde response gene expression from RNA-seq samples collected from same experiment as Fig 3A. (B) Venn diagrams representing overlap between genes induced or repressed in FNP^QiI^ treatment and MNR. (C) RT-qPCR analysis of *bfr1*, *obr1*, *pmd1* and *nde1* expression in untreated cells (UT) compared to cells exposed to FNP^QiI^ for 24 hours (0 hours) or cells given 6-48 hours recovery in non-selective media following FNP^QiI^ washout. Significance: *Student’s t-test; *p<0.05, **p<0.01*, *\*\*\*<0.001, ****<0.0001*. (D) Efflux of Rhodamine 6G (R6G) in WT or *4xpump*Δ cells untreated (UT) or treated with FNP^QiI^. with (solid lines) or without glucose (dashed lines). Efflux of R6G is plotted over time.

**S3 Fig.**
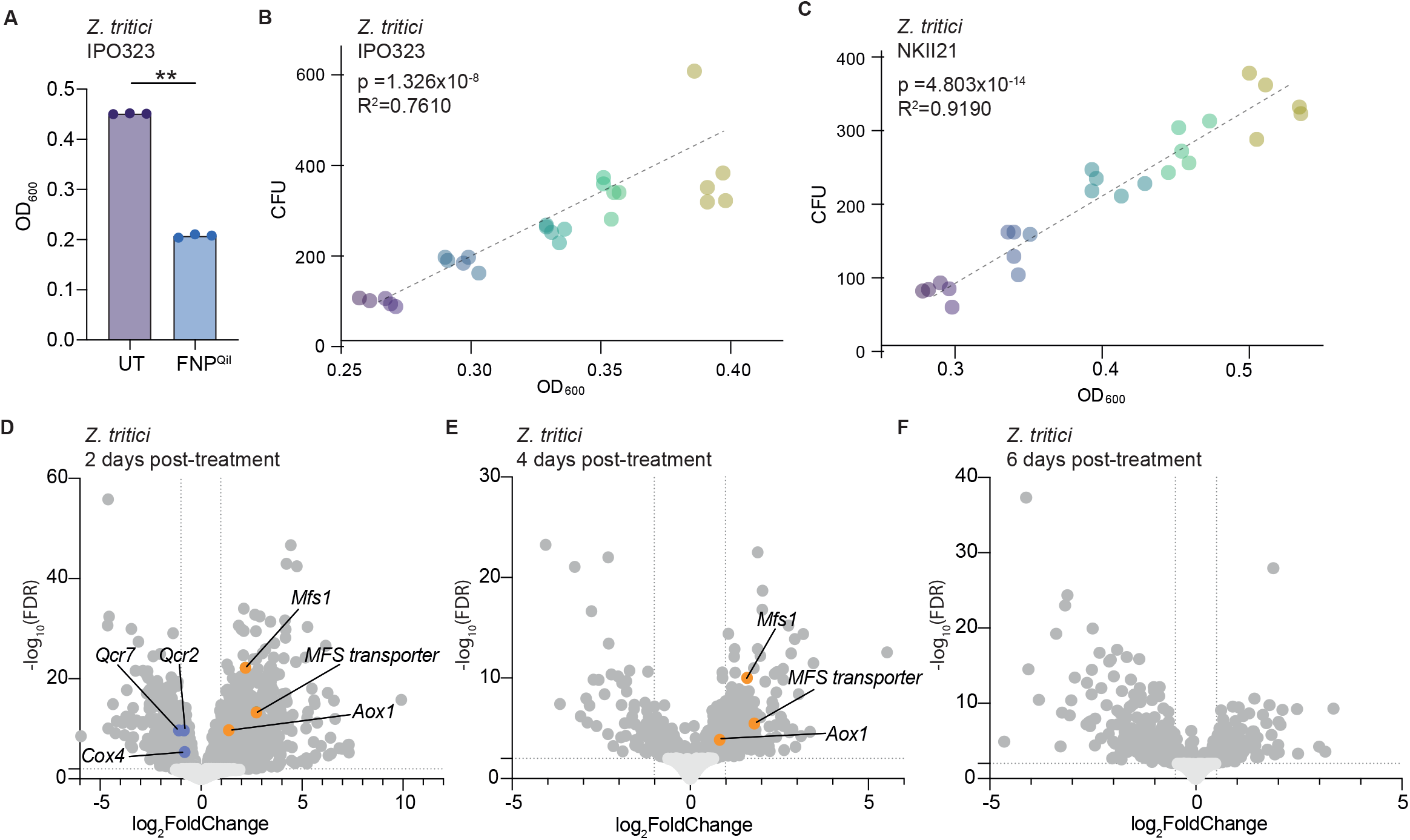
FNP^QiI^ exposure increases resistance of *Z. tritici* to azole^DMI^-based antifungals. (A) *Z. tritici* IPO323 cells were grown in YPD only (<u>U</u>n<u>T</u>reated) or with addition of FNP^QiI^. Cell growth was measured by optical density OD_600_. (B) Microwell plate liquid growth assays using IPO323 showing OD_600_ comparing untreated (UT) cells and those grown for 24 hours in FNP^QiI^ followed by washout and non-selective growth for indicated time (hours) (as in Fig 4A). (C) Microwell plate liquid growth assays using NKII21 showing OD_600_ comparing untreated (UT) cells and those grown for 24 hours in FNP^QiI^ followed by washout and non-selective growth for indicated time (hours) (as in Fig 4A). (D-F) Volcano plots from RNA-seq of in *Z. tritici* IPO323 following 24h FNP^QiI^ treatment with 2 (D), 4 (E) or 6 (F) days recovery in YPD after washout, samples compared to untreated cells. Genes with p-value <0.01 and fold change value ≥1.5 are indicated (dark grey). Representative DEGs are indicated in orange (up) and purple (down).

**Supplementary Table 1.**
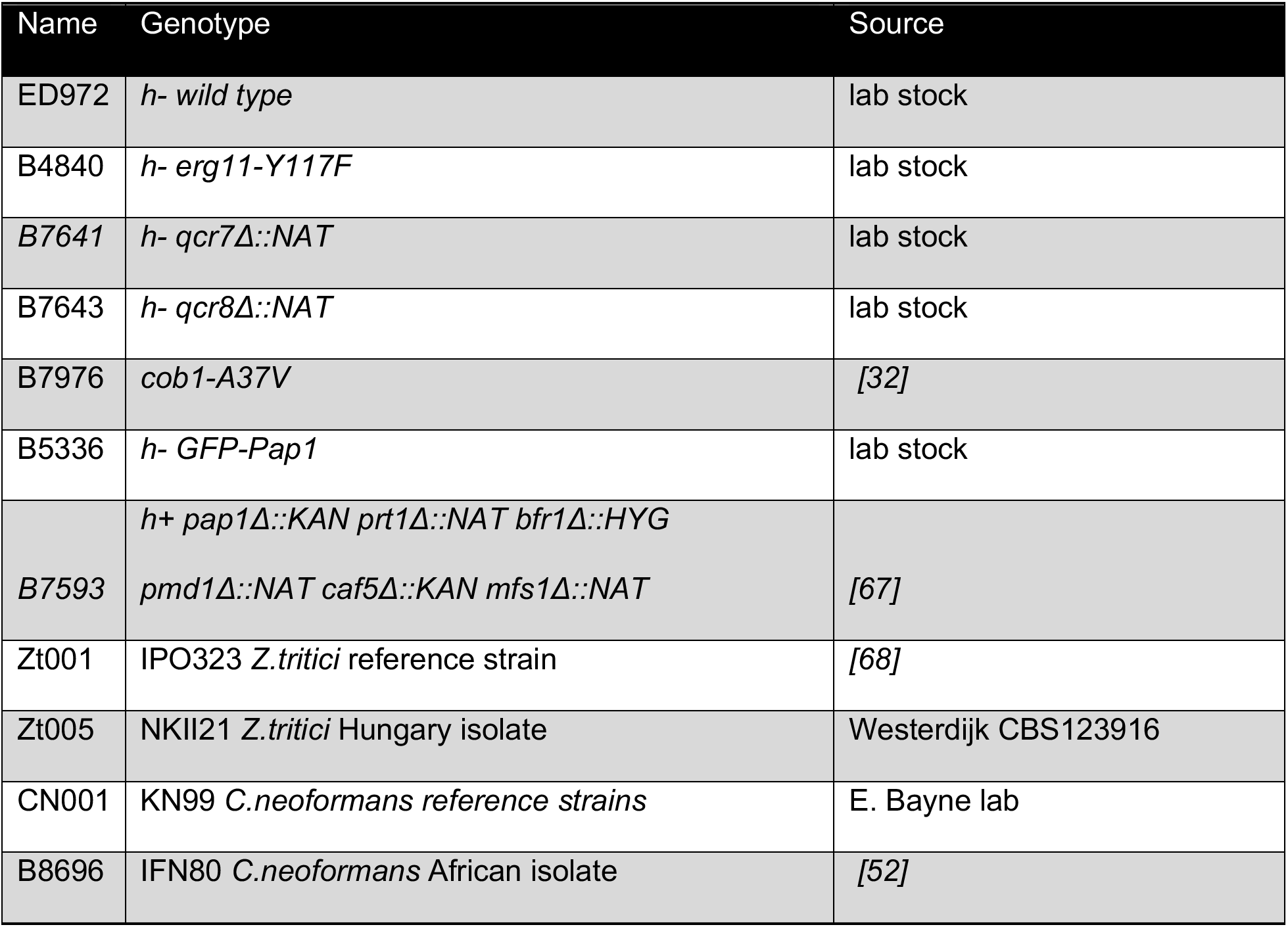
Strains used in this study.

| Name | Genotype | Source |
|---|---|---|
| ED972 | *h- wild type* | lab stock |
| B4840 | *h- erg11-Y117F* | lab stock |
| *B7641* | *h- qcr7Δ::NAT* | lab stock |
| B7643 | *h- qcr8Δ::NAT* | lab stock |
| B7976 | *cob1-A37V* | *[32]* |
| B5336 | *h- GFP-Pap1* | lab stock |
| *B7593* | *h+ pap1Δ::KAN prt1Δ::NAT bfr1Δ::HYG*<br>*pmd1Δ::NAT caf5Δ::KAN mfs1Δ::NAT* | *[67]* |
| Zt001 | IPO323 *Z.tritici* reference strain | *[68]* |
| Zt005 | NKII21 *Z.tritici* Hungary isolate | Westerdijk CBS123916 |
| CN001 | KN99 *C.neoformans reference strains* | E. Bayne lab |
| B8696 | IFN80 *C.neoformans* African isolate | *[52]* |

**Supplementary Table 2.**
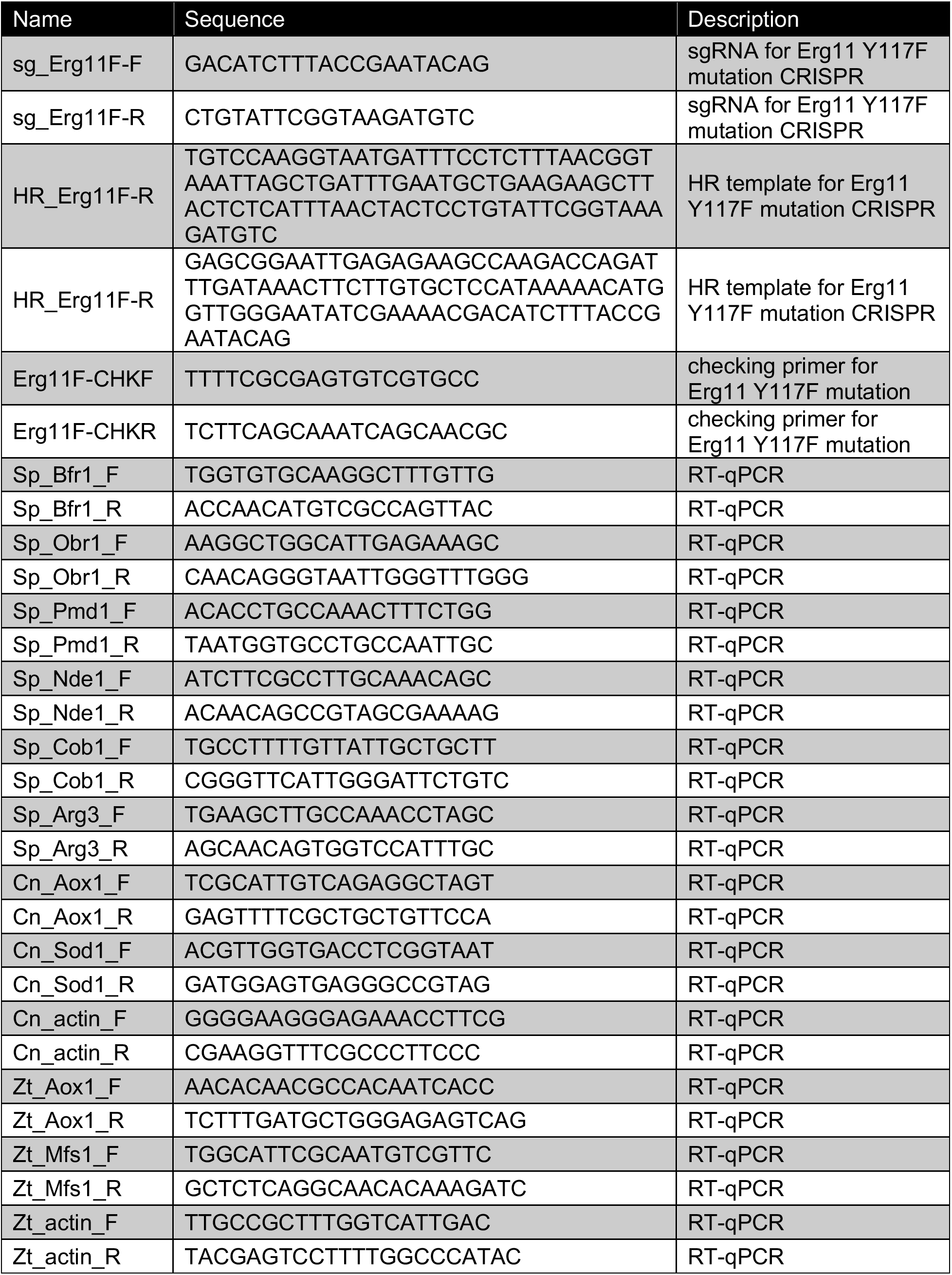
Oligonucleotides used in this study.

